# An improved 3DMax algorithm to reconstruct the three-dimensional structure of the chromosome

**DOI:** 10.1101/2020.07.09.195693

**Authors:** Liwei Liu, Huili Yao

## Abstract

In recent years, with the development of high-throughput chromosome conformation capture (Hi-C) technology and the reduction of high-throughput sequencing cost, the data volume of whole-genome interaction has increased rapidly, and the resolution of interaction map keeps improving. Great progress has been made in the research of 3D structure modeling of chromosomes and genomes. Several methods have been proposed to construct the chromosome structure from chromosome conformation capture data. Based on the Hi-C data, this paper analyses the relevant literature of chromosome 3D structure reconstruction and it summarizes the principle of 3DMAX, which is a classical algorithm to construct the 3D structure of a chromosome. In this paper, we introduce a new gradient ascent optimization algorithm called XNadam that is a variant of Nadam optimization method. When XNadam is applied to 3DMax algorithm, the performance of 3DMax algorithm can be improved, which can be used to predict the three-dimensional structure of a chromosome.

**Author summary:** The exploration of the three-dimensional structure of chromosomes has gradually become a necessary means to understand the relationship between genome function and gene regulation. An important problem in the construction of three-dimensional model is how to use the interaction map. Usually, the interaction frequency can be transformed into the spatial distance according to the deterministic or non-deterministic function relationship, and the interaction frequency can be weighted as weight in the objective function of the optimization problem. When the frequency of interaction is weighted as weight in the objective function of the optimization problem, what kind of optimization method is used to optimize the objective function is the problem we consider. In order to solve this problem, we provide an improved stochastic gradient ascent optimization algorithm(XNadam). The XNadam optimization algorithm combined with maximum likelihood algorithm is applied to high resolution Hi-C data set to infer 3D chromosome structure.

## Introduction

The 3D structure of chromosomes is critical for the study of DNA replication, gene regulation, genome interaction, genome folding, and genome function. How to predict the 3D structures of a chromosome by using computational technology and bioinformatics has become the core of genomics research[1-9]. At present, according to the different principles of constructing chromosome 3D structures, the structural model can be divided into two categories: probability restraint-based prediction model and distance restraint-based prediction model. Because the 3D structure of chromosomes in cells is dynamic, we can use some probability distribution to describe it, so we can transform the problem of constructing 3D structure of chromosomes into the problem of building probability model. Methods that implement this process are known as probability restraint-based prediction model. Some of these methods are based on a two-step process to perform 3D structural modeling of the chromosome genome, which involves converting the interaction frequency (IF) between the pair of fragments in the Hi-C data to the distance between them, and then optimizing the distance value and inferring the best 3D structure meets the distance. The method to achieve this two-step process is called a distance restraint-based model. Several of such methods have been proposed, each of which varies in restraint representation and optimization methods adopted. In the process of distance optimization, a gradient iterative optimization algorithm is often used to optimize the objective function. At present, SGD, Momentum, Nesterov, Adagrad, Adadelta, Adam, Adamax and Nadam[10-16] are the existing stochastic gradient optimization algorithms. In this paper, according to the advantages and disadvantages of the existing stochastic gradient optimization algorithm, we introduce a new Gradient ascent optimization algorithm called XNadam that is a variant of Nadam optimization method.

## Materials and methods

### 3DMax methods

3DMax methods[17] is distance restraint-based model proposed by Oluwatosin et al. It combines a maximum likelihood algorithm and a gradient ascent method to generate optimized structures for chromosomes. 3DMax structure modeling starts with a random initialization of the coordinates of all the regions, and then transforms the contact matrix into space distance. By assuming that each data point (interaction frequencies or distances) are conditionally independent, it obeys the normal distribution. So as to construct a log likelihood function as an objective function. Then, the gradient ascent method is used to iteratively optimize the objective function until the algorithm converges, and the chromosome 3D coordinates are updated synchronously during the iteration.

Oluwatosin et al. also implemented a variant of the 3DMax algorithm above, called 3DMax1. In 3DMax1 structure modeling, it uses a stochastic gradient ascent algorithm which is called the adaptive Gradient algorithm (AdaGrad) to optimize the log likelihood objective function. Adagrad [18] is a gradient-based optimization algorithm, which optimizes the iteration of the program by dividing the learning rate by the square root of the sum of the squares of the historical gradients. It performs larger updates for infrequent or sparse parameters and smaller updates for frequent or fewer sparse parameters. Therefore, when dealing with sparse parameters, it usually has better convergence performance than some other stochastic gradient ascent algorithm. Its disadvantage is that the denominator will continue to accumulate and the memory requirement is large, so the learning rate will shrink and eventually become very small, or even zero.

### XNadam methods

At present, SGD, Momentum, Nesterov, Adagrad, Adadelta, Adam, Adamax and Nadam are the existing stochastic gradient optimization algorithms. In this paper, we introduce a new Gradient ascent optimization algorithm called XNadam that is a variant of Nadam optimization method. Nadam (Nesterov-accelerated Adaptive Moment Estimation) is a combination of Adam and Nesterov algorithm, with the advantages of both. It has a stronger constraint on the learning rate, which makes the algorithm better in some problems. The main advantage of Nadam is that each iterative learning rate has a certain range, and different adaptive learning rates are calculated for different parameters, after bias correction. Nadam makes the parameters more stable, while the memory demand is small, and it is also suitable for most non-convex optimization, big data and high dimensional space. Due to the addition of Nesterov term, a correction is made during gradient updating to avoid too fast forward and improve sensitivity. XNadam that is a variant of Nadam optimization method and its principle is similar to Nadam. The difference is that XNadam decay the learning rate with a sinusoidal correlation method for each batch, and also uses sinusoidal correlation method to correct the first-order moment estimation. Therefore, XNadam has all the advantages of Nadam algorithm. At the same time, it will be stronger for learning rate constraint and for data, it performs better than most other optimization methods. XNadam algorithm is described in detail in Table 1.

**Table 1.**
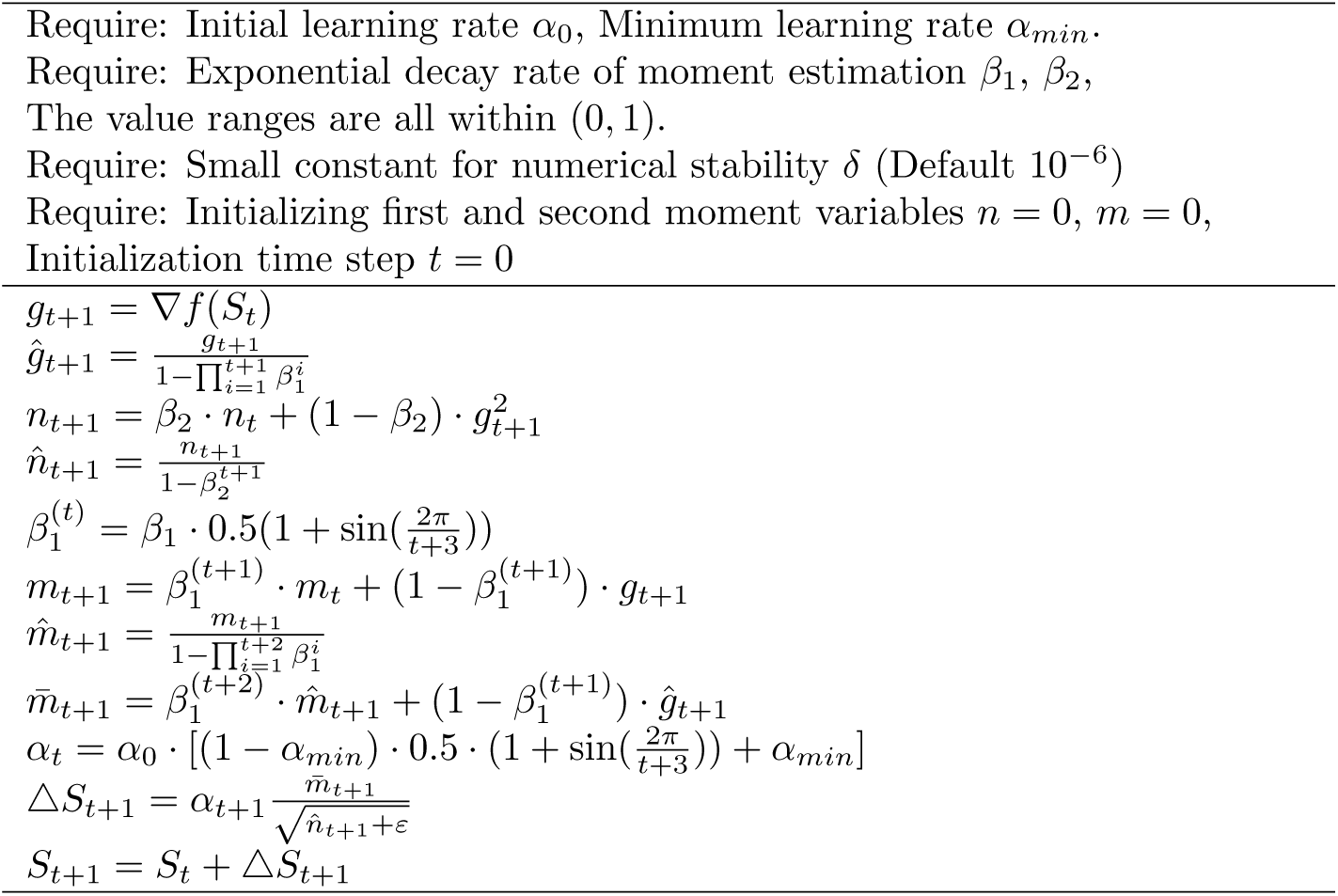
XNadam Algorithm

### Measurement of model similarity and accuracy

To test this algorithm, we compared the performance of 3DMaxAdaGrad, 3DMaxNadam and 3DMaxXNadam. 3DMaxAdaGrad, 3DMaxNadam and 3DMaxXNadam are variants of 3DMax algorithm, among which 3DMaxAdaGrad is a prediction model combining a maximum likelihood algorithm and AdaGrad algorithm to reconstruct the 3D structure of a chromosome, while 3DMaxNadam is a prediction model combining a maximum likelihood algorithm and Nadam optimization algorithm, 3DMaxXNadam is a prediction model combining a maximum likelihood algorithm and XNadam optimization algorithm. 3DMaxXNadam algorithms used *β*_1_ = 0.6, *β*_2_ = 0.9999, *α*_*min*_ = 0.0001. Only the learning rate across differed across algorithms and experiments.

We used the Distance Pearson Correlation Coefficient (DPCC), the Distance Spearman Correlation Coefficient (DSCC), the Distance Root Mean Square error (DRMSE)[19-22], Kullback–Leibler divergence, Cross Entropy to measure the similarities between chromosomal structures, and assess the accuracy of the constructed structures as in the previous studies.

In this study, the Hi-C contact frequency data of the yeast 1-12 chromosome were normalized using the Yaffe and Tanay [23]normalization technique was adopted. Table 2 shows the number of the chromosomal locus.

**Table 2.**
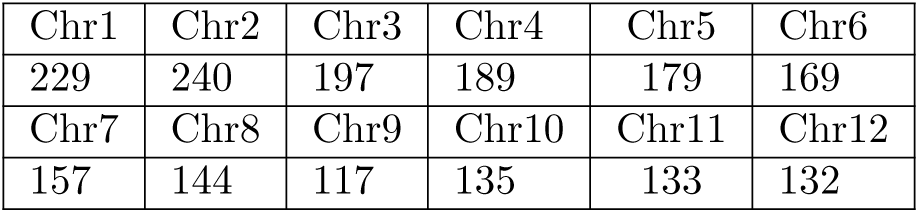
the number of the chromosomal locus

| Chr1 | Chr2 | Chr3 | Chr4 | Chr5 | Chr6 |
| --- | --- | --- | --- | --- | --- |
| 229 | 240 | 197 | 189 | 179 | 169 |
| Chr7 | Chr8 | Chr9 | Chr10 | Chr11 | Chr12 |
| 157 | 144 | 117 | 135 | 133 | 132 |

## Results

The average DSCCDPCC and DRMSE value between 3DMaxAdaGrad, 3DMaxNadam, 3DMaxXNadam algorithm and the distance matrix for 1-12 contact matrices respectively, the results are shown in Fig 1–Fig 4.

**Fig 1.**
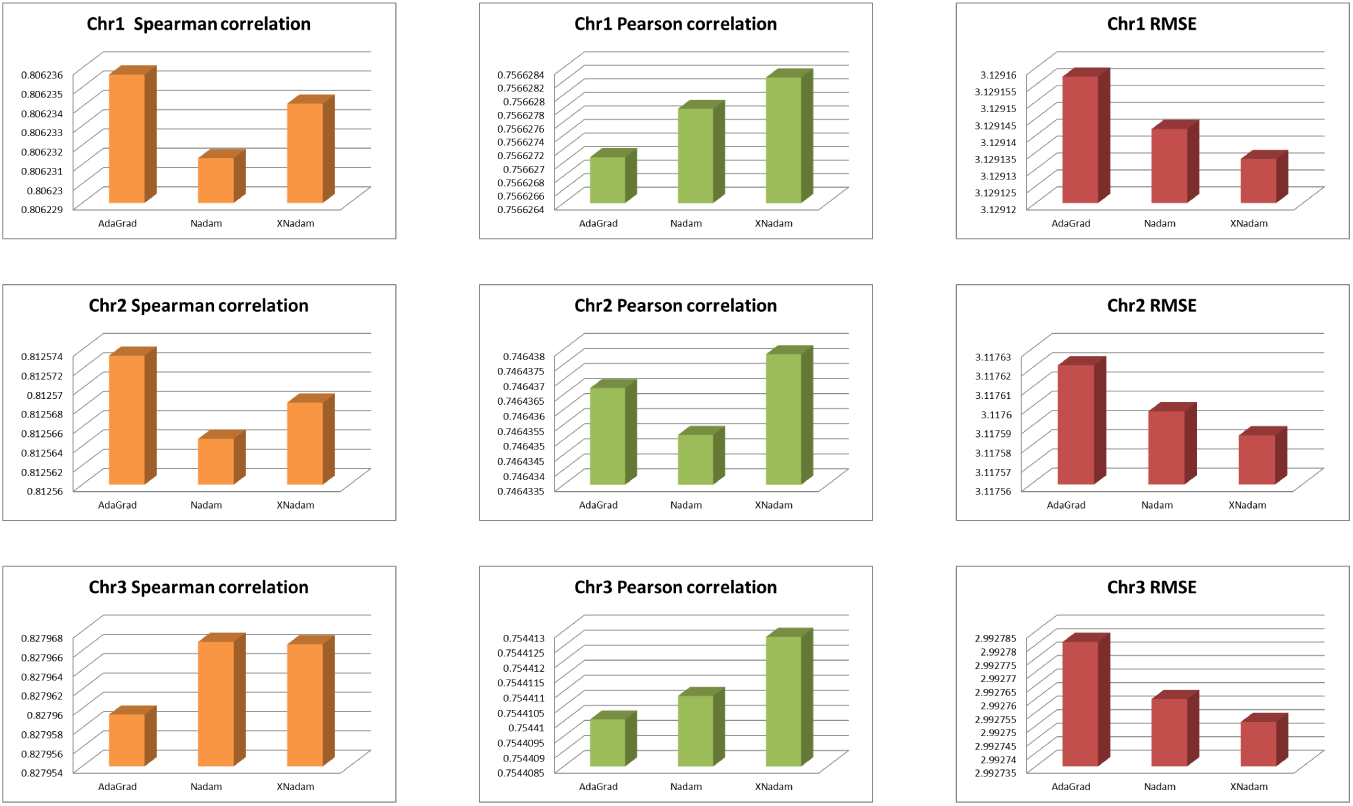
The evaluation results for 3DMaxAdaGrad, 3DMaxNadam and 3DMaxXNadam(Ch1-Ch3)

**Fig 2.**
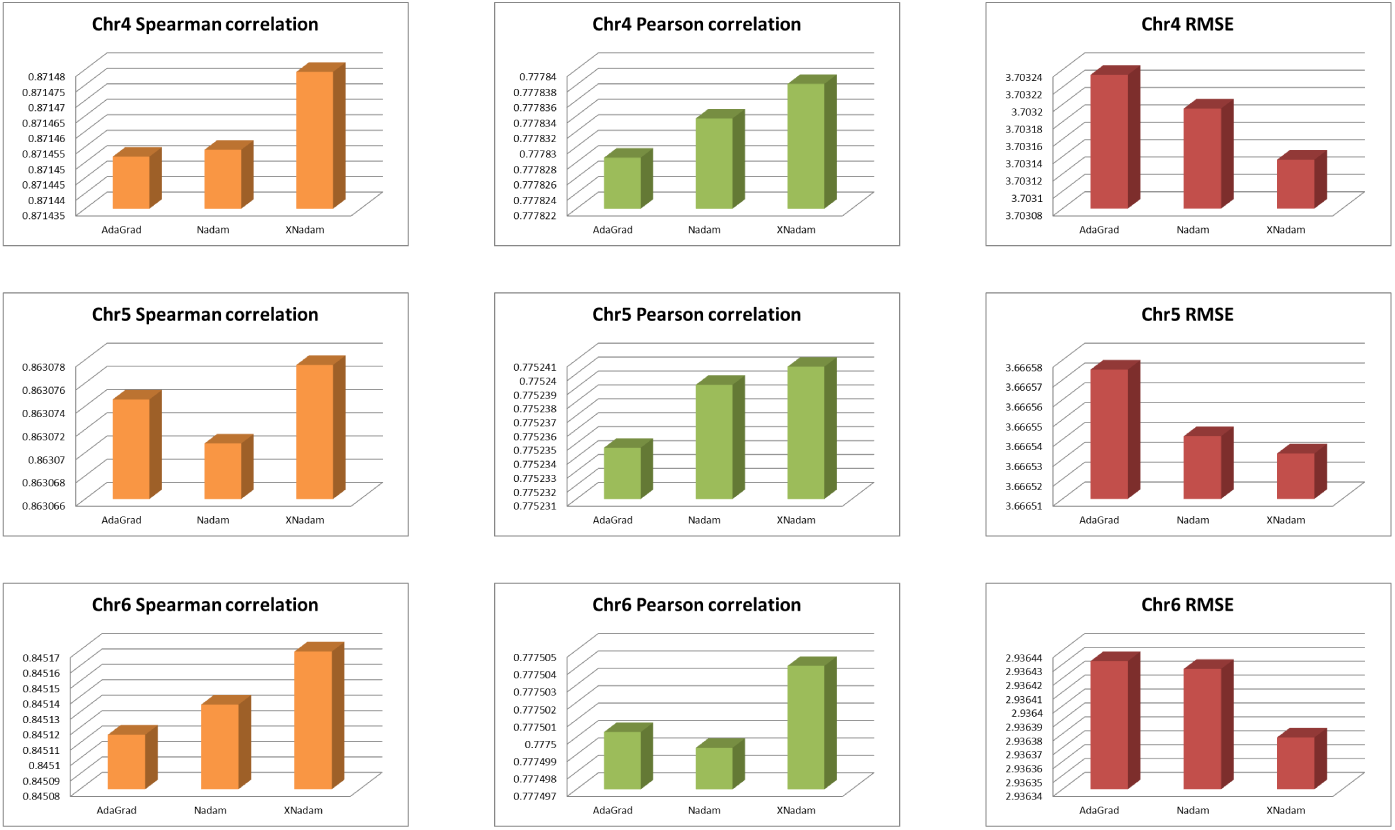
The evaluation results for 3DMaxAdaGrad, 3DMaxNadam and 3DMaxXNadam(Ch4-Ch6)

**Fig 3.**
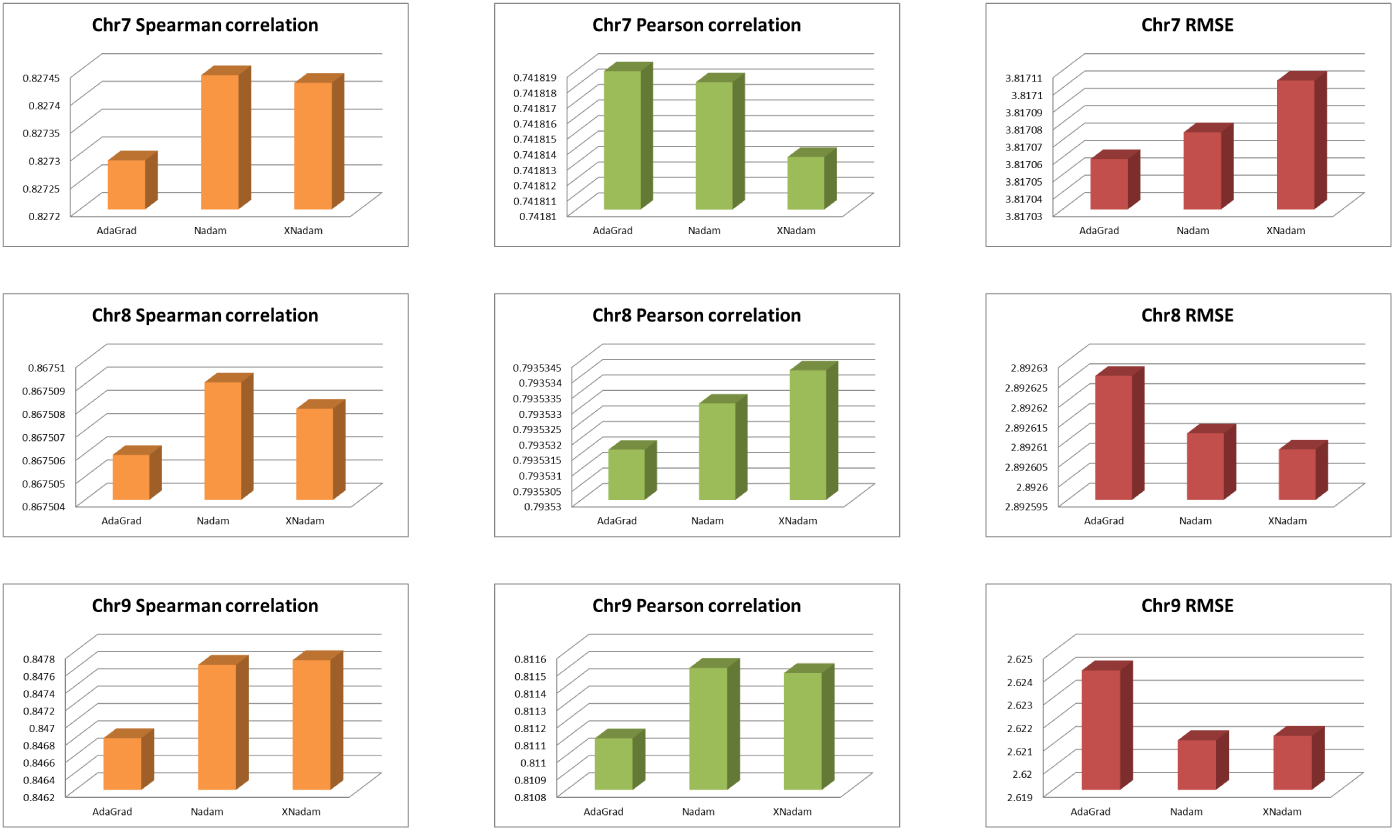
The evaluation results for 3DMaxAdaGrad, 3DMaxNadam and 3DMaxXNadam(Ch7-Ch9)

**Fig 4.**
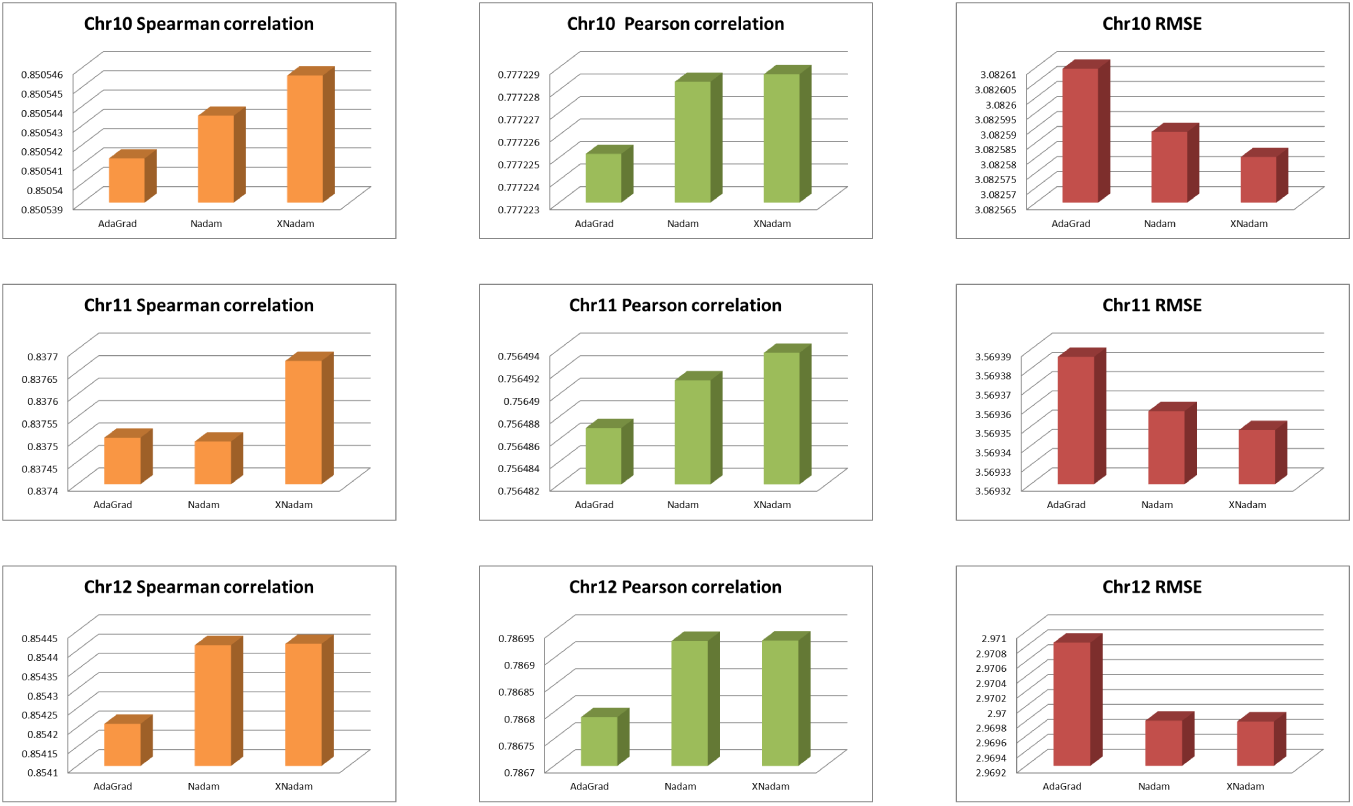
The evaluation results for 3DMaxAdaGrad, 3DMaxNadam and 3DMaxXNadam(Ch10-Ch12)

## Discussion

The histogram above shows the DSCC, DPCC and DRMSE evaluation values of 3DMaxAdaGrad, 3DMaxNadam and 3DMaxXNadam algorithm to reconstruct the 3D chromosome structure. The first column of the yellow histogram represents the DSCC value of 3DMaxAdaGrad, 3DMaxNadam and 3DMaxXNadam algorithm for reconstructing the 3D structure of 1-12 chromosomes. It can be seen from the observation figures that for chromosomes 1 and 2, the DSCC value of 3DMaxAdaGrad algorithm for reconstructing the 3D structure of chromosomes is the largest, followed the DSCC value of 3DMaxXNadam algorithm. The DSCC value of 3DMaxnadam algorithm is the smallest. When observing chromosome 8, the DSCC value of 3DMaxNadam algorithm for reconstructing the 3D structure of chromosomes is the largest, that of 3DMaxXNadam algorithm is the second largest, and the DSCC value of 3DMaxAdaGrad algorithm is the smallest. In addition to the three chromosomes mentioned above, the DSCC value of all the other chromosomes is the highest in the 3DMaxXNadam algorithm.

The second column of the green histogram represents DPCC values of 3DMaxAdaGrad,3DMaxNadam and 3DMaxXNadam algorithm for reconstructing the 3D structure of 1-12 chromosomes. It can be seen from the figure that in addition to chromosome 7, among the other chromosomes, the DPCC value of the 3DMaxXNadam algorithm is the largest, the DPCC value of the 3DMaxNadam algorithm is the second, and the DPCC value of the 3DMaxAdaGrad algorithm is the smallest.

The third column of the red histogram represents DRMSE values of 3DMaxAdaGrad,3DMaxNadam and 3DMaxXNadam algorithm for reconstructing the 3D structure of 1-12 chromosomes. It can be seen from the figure that in addition to chromosome 7, among the other chromosomes, the DRMSE value of the 3DMaxXNadam algorithm is the smallest, the DRMSE value of the 3DMaxNadam algorithm is the second, and the DRMSE value of the 3DMaxAdaGrad algorithm is the largest.

According to the above experimental results, the performance of 3DMaxNadam is better.

Compared with existing algorithms, the 3DMaxXNadam algorithm has high accuracy in the reconstruction of chromosome 3D structure. In this paper, the algorithm is used to reconstruct the 3D spatial structure of 1-12 chromosomes, and the 3D effect map of 1-12 chromosomes is obtained (as shown in Fig 5). Fig 5 shows the 3D effect map of chromosomes, which shows the coiling and folding shape of chromosome in space.

**Fig 5.**
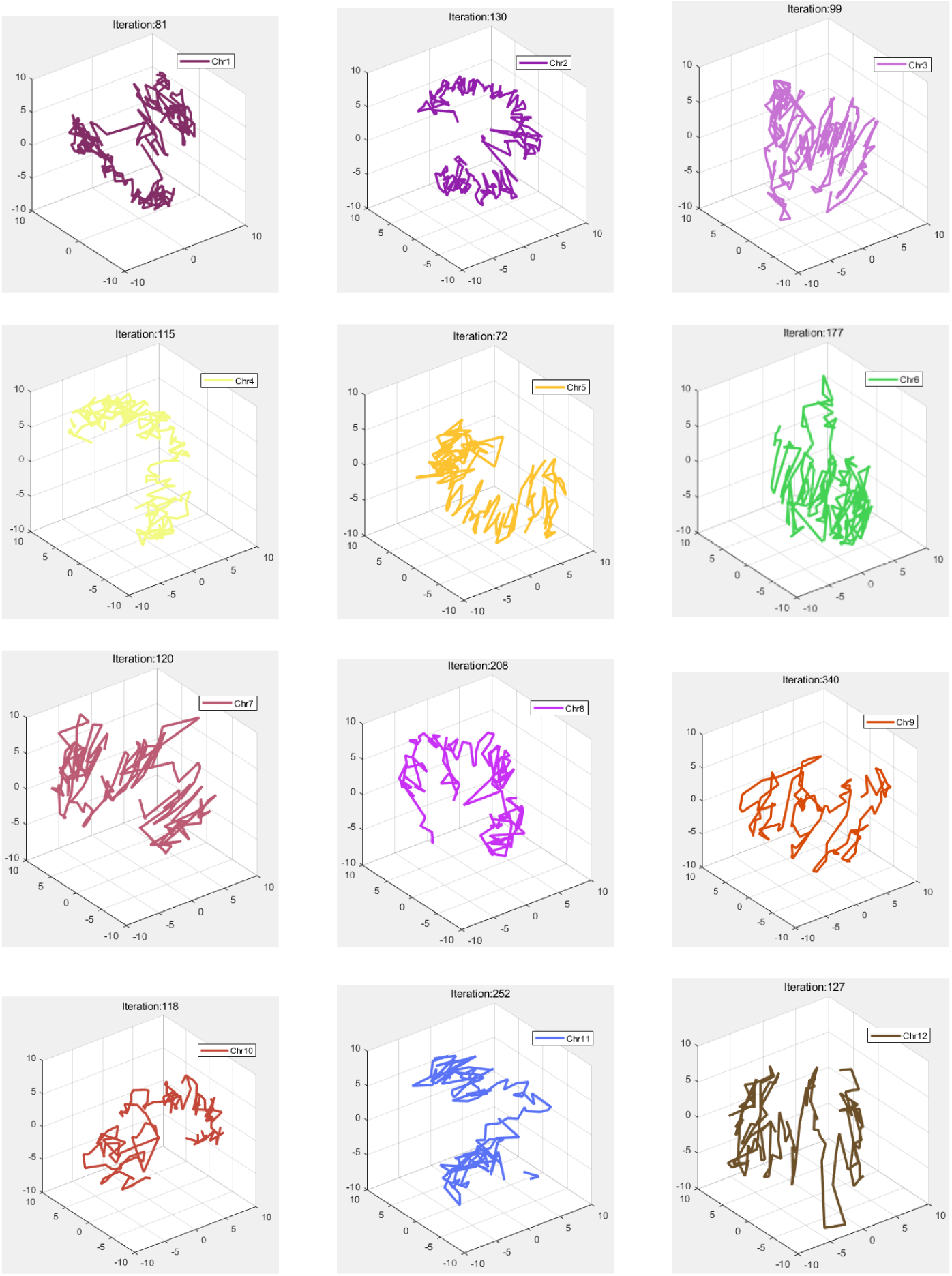
The 3D effect map of chromosomes

## Conclusion

In this paper, we developed a new gradient ascent optimization algorithm (XNadam) based on Nadam, and we introduce 3DMaxAdaGrad, 3DMaxNadam and 3DMaxXNadam algorithm. 3DMaxAdaGrad, 3DMaxNadam and 3DMaxXNadam algorithm are used to reconstruct the three-dimensional structure of yeast chromosome, and the values of DSCC, DPCC and drmse of the generated optimized structure of chromosome are calculated to measure the performance of the prediction method. The results on synthetic datasets show that 3DMaxXNadam of combining a maximum likelihood algorithm and XNadam algorithm can effectively reconstruct chromosomal models from Hi-C contact matrices normalized, at the same time, it is faster and has a low memory requirement compared to some other methods. This conclusion shows that when the new gradient ascent optimization algorithm (XNadam) optimizes the log likelihood objective function, it performs better than most other optimization methods.

## Acknowledgments

This research was supported by China Postdoctoral Science Foundation (Grant No. 2018M631782), the Foundation of Liaoning Educational Committee (Grant No. JDL2019032), and the Natural Science Foundation of Liaoning Province of China (Grant No. 201800278).

